# A brain signature of vaping cue-elicited craving

**DOI:** 10.64898/2026.10.15.759290

**Authors:** Shangcheng Zhao, Hongbo Yu, Leonie Koban, Yidi Wang, Qiyao Peng, Tianlai Ye, Michael Amlung, James MacKillop, Lawrence H. Sweet, Jiaying Liu

**Author notes:** Equal contribution (Shangcheng Zhao, Hongbo Yu). Correspondence should address to: Jiaying Liu, Ph.D., Department of Communication, 4425 Social Sciences and Media Studies Building, University of California Santa Barbara, Santa Barbara, CA 93106-4020., Lawrence H. Sweet, Ph.D. Department of Psychology, Department of Psychology, 159 Psychology Building, 125 Baldwin Street, University of Georgia, Athens, GA 30602.

## Abstract

E-cigarette use among young adults has grown rapidly, raising concerns about nicotine dependence and motivating a better understanding of the neurobiology of vaping. We used a cue-induced craving fMRI paradigm to develop and evaluate a brain signature of vaping cue-elicited craving, and to compare it with smoking– and food-craving signatures developed using the same analytic pipeline. Using LASSO–PCR with leave-one-subject-out cross-validation, we identified a distributed pattern of brain activity that reliably predicted normative vaping craving from an independent sample (hereafter, normative craving). The vaping-craving signature was also substantially distinct from smoking– and food-craving signatures, both spatially and functionally, and showed specificity relative to previously established craving signatures. Moreover, greater signature expression was associated with greater e-cigarette dependence severity and showed a positive, though modest, prospective association with daily puff counts over four weeks, demonstrating the signature’s behavioral and clinical relevance. Together, these findings establish a cross-validated brain signature of vaping cue-elicited craving, demonstrate its differentiation from other craving-related neural patterns, and link its expression to e-cigarette dependence and real-world use. More broadly, they support the potential of this framework for developing reproducible neural markers of craving.

**Significant statement:** Craving is a basic human experience, but also a powerful engine for addiction. Although vaping has become a common form of nicotine use, we still know little about vaping-specific craving. Craving is currently measured mostly through self-report, which is prone to bias. Using neuroimaging and machine learning, we identified a brain signature that tracks e-cigarette dependence and predicts cue-related craving tendencies. This signature captures vaping-specific craving above and beyond a previously established domain-general signature of craving, and is distinct from brain-based signatures of food and cigarette craving. These findings highlight a distinct neural representation for vaping craving and provide a new and generalizable brain-based measure for predicting vaping dependence.

## Introduction

Electronic cigarette (e-cigarette) use, commonly known as vaping, has increased rapidly over the past decade. Although vaping may serve as a harm-reduction tool for previous cigarette smokers, its widespread use among non-smokers, particularly among teenagers and young adults, has raised concerns about nicotine initiation and dependence. National surveillance data show that e-cigarettes have a higher prevalence among young adults than traditional cigarettes. In 2024, 14.8% of U.S. adults aged 18–24 reported using e-cigarettes, compared to 3.4% who reported using traditional cigarettes (Sohi & Ng, 2026). Regular vaping can lead to nicotine dependence and has also been associated with subsequent combustible cigarette smoking (Chan et al., 2021; Leventhal et al., 2016). Because craving and nicotine dependence are central features of problematic e-cigarette use, developing measures that complement self-report may improve the characterization of vaping-related craving and dependence. However, current assessments rely heavily on self-reported measures, which are susceptible to reporting biases and may not fully capture the neural processes associated with cue-elicited craving.

One promising approach is to measure cue reactivity, a hallmark of substance use disorders. Cue reactivity arises through classical conditioning, whereby repeated pairing of drug consumption with environmental stimuli causes previously neutral cues to acquire motivational value (Stewart et al., 1984). As a result, substance-related objects, contexts, advertisements, and other environmental cues can elicit conditioned responses even in the absence of drug administration. Extensive evidence shows that exposure to substance-associated cues reliably evokes subjective craving (Betts et al., 2021; Carter & Tiffany, 1999), and can be accompanied by dopamine release and altered activity in brain regions involved in reward and salience processing, including the ventral striatum (Chase et al., 2011; Kühn & Gallinat, 2011; Volkow et al., 2004). Importantly, cue-induced neural responses predict subsequent drug-seeking behavior, including lapse and relapse during cessation, supporting the relevance of cue reactivity to addiction-related behavior and subsequent substance use (Owens et al., 2018; Suzuki & Kober, 2018).

Functional magnetic resonance imaging (fMRI) combined with cue-reactivity paradigms provides a powerful method for characterizing the brain mechanisms underlying craving. Several pioneering studies have examined neural responses to e-cigarette-related cues. These studies consistently reported increased activation in brain regions involved in reward processing, including the nucleus accumbens, as well as regions supporting cognitive control, such as the middle frontal gyrus (Chen et al., 2018; Garrison et al., 2018; Grundinger et al., 2025; Liu et al., 2025). Together, these findings suggest that vaping cues engage neural systems implicated in reward, motivational salience, and cognitive control. However, these studies relied primarily on ROI-based or univariate whole-brain analyses to characterize condition-related differences in regional activation. While informative for identifying neural systems responsive to vaping-related stimuli, these approaches do not address the complementary predictive question of whether distributed patterns of brain activity can be used to estimate the craving intensity associated with particular cues across participants. A multivariate predictive approach therefore shifts the emphasis from identifying condition-related brain responses toward out-of-sample brain-to-craving prediction.

Recent work combining fMRI with multivariate machine learning has demonstrated that many psychological experiences, such as physical pain, reward, and negative affect, can be characterized using distributed patterns of whole-brain activity, often referred to as *brain signatures* (Čeko et al., 2022; Speer et al., 2023; Wager et al., 2013). Unlike conventional univariate analyses, brain signatures predict psychological states directly from distributed patterns of brain activity, enabling brain-to-behavior and brain-to-experience prediction at the individual level (Woo et al., 2017; Krage, et al., 2018). This approach offers several advantages for addiction research. First, it provides sensitive, brain-based predictive measures that can complement conventional self-report assessments of craving. Second, it helps characterize distributed neural representations associated with craving, potentially facilitating the development of biologically informed interventions (Venniro et al., 2020). Finally, multivariate signatures can be evaluated directly in held-out participants and independent datasets. Such out-of-sample testing does not by itself establish a reproducible biomarker, but it provides an important test of whether a predictive pattern transfers beyond the training data (Woo et al., 2017). Several multivariate brain signatures of craving have now been reported across substance-use domains (Kulkarni et al., 2023; Mahdavi-Doost et al., 2025; Tian et al., 2024). Most directly relevant to the present study, Koban et al. (2023) developed a neurobiological craving signature (NCS) that predicted food and drug craving, including craving for combustible cigarettes, alcohol, and cocaine. E-cigarette cues were not included in its development, leaving open whether vaping cue-elicited craving is adequately captured by a broader, domain-general craving representation or whether it is better characterized by a vaping-specific predictive pattern.

In the present study, we developed and evaluated a whole-brain brain signature of vaping cue-elicited craving using fMRI and multivariate machine learning. An independent sample of 442 young adult regular vapers rated vaping, smoking, and food images, providing image-level normative craving ratings. A separate fMRI sample of 66 young adult regular vapers who reported no combustible cigarette smoking in the past 30 days viewed these images during scanning. We first developed and cross-validated a whole-brain brain signature predicting normative vaping-cue craving intensity. To evaluate its specificity, we developed corresponding smoking– and food-craving signatures, examined cross-cue predictive transfer, and compared the vaping signature with the previously established NCS. We also examined whether vaping-signature expression was associated with e-cigarette dependence and subsequent vaping behavior. Finally, as a proof-of-principle test of cross-dataset portability, we applied the smoking signature developed using the same analytic pipeline to an independent smoking-cue fMRI dataset and tested whether vaping signature showed corresponding transfer to indicate specificity.

We expected the vaping model to predict normative vaping-cue craving intensity within the primary sample and to show limited transfer to smoking and food cues. As a proof-of-principle cross-dataset test, we further expected the smoking signature to distinguish smoking from neutral cues in the independent smoking dataset, whereas the vaping signature was expected to show limited transfer to smoking cues.

## Results

### Description of Brain Signatures

We developed brain signatures that predicted continuous, image-specific craving intensity (aggregated rating from an independent sample) from fMRI contrast images using least absolute shrinkage and selection operator–principal component regression (LASSO–PCR) (Koban et al., 2023) with leave-one-subject-out cross-validation. Category-specific cues were categorized into five craving levels (1–5) based on group-average ratings, and individual brain activation maps were estimated using first-level GLMs. Separate signatures were subsequently trained for vaping, smoking, and food craving intensity from those category-specific brain maps. Each signature comprises a voxel-wise whole-brain weight map and an intercept: within the full multivariate pattern, positive weights contribute to higher pattern expression, whereas negative weights contribute to lower pattern expression. Because these weights are conditional on all other features in the model, they should not be interpreted as independent regional activation effects. Models were fit on all but one participant per fold and evaluated on held-out data.

When applied to a test image, a signature returns one pattern-expression score, which is computed by adding the intercept to the dot product of the test image with the weight map. This score represents the model-predicted normative craving level associated with the cue-evoked brain pattern.

Figure 1A, 2A, and 3A show thresholded weight maps from bootstrapping, while the predictions and pattern expression are calculated based on the full unthresholded map; the thresholded map highlights regions with the most reliable positive and negative contributions. For the vaping signature, bootstrap-stable positive weights were observed in bilateral occipital poles and sensorimotor cortex, while bootstrap-stable negative weights were observed in the right occipital cortex, right anterior cingulate cortex, and left thalamus. For the smoking signature, positive weights were observed in bilateral occipital cortex and left superior/middle frontal gyrus, while negative weights were found in bilateral occipital poles, right precentral gyrus, and right primary motor cortex. For the food signature, positive weights were observed in right lateral occipital cortex and left occipital pole, while negative weights were found in left lingual gyrus, left nucleus accumbens, right occipital pole, and right precentral gyrus.

**Figure 1.**
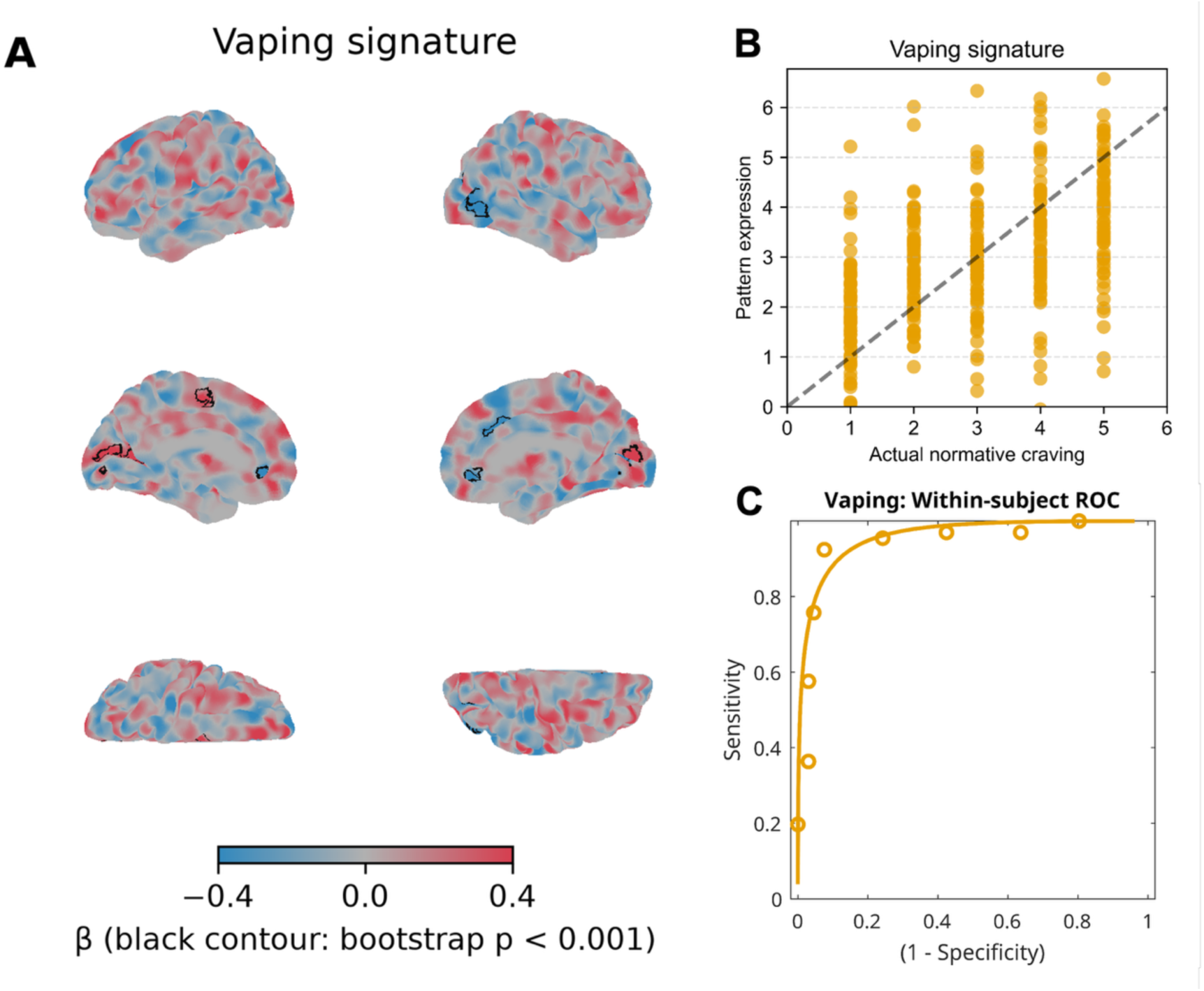
Vaping craving brain signature. **A.** Medial, lateral, and axial views of the brain signature weights. Red and blue indicate positive and negative LASSO weights, respectively, with color saturation reflecting the absolute magnitude of the weights. Regions enclosed by black contours survived bootstrap testing at *p* < .001. **B.** Vaping brain signature pattern expression versus normative craving ratings. **C.** Within-subject receiver operating characteristic (ROC) curve for discriminating the highest versus lowest levels of craving.

### Predictive Performance of Brain Signatures

We evaluated each brain signature’s predictive performance by fitting a Multiple Linear Regression model to examine the association between brain signature pattern expression and normative craving, controlling for sex, age, body mass index (BMI), and average head motion. Normative craving rating was the dependent variable and pattern expression value was the independent variable, and cluster-robust standard errors were estimated at the participant level to account for within-subject dependence.

Across all cue types, pattern expression scores remained positively associated with normative craving level after controlling for covariates (vaping: *β* = 0.52 ± 0.05, *t*(61) = 9.77, *p* < .001; smoking: *β* = 0.60 ± 0.03, *t*(61) = 17.97, *p* < .001; food: *β* = 0.66 ± 0.03, *t*(61) = 19.43, *p* < .001). Linear mixed models with a random intercept of subject and a random slope of pattern expression by subject were fitted but did not outperform the simpler multiple regression model with subject-clustered standard errors (See Supplementary Results). As a further validation using participants’ own self-reported craving ratings collected during scanning, vaping-signature pattern expression showed a positive, though not statistically significant, association with block-wise vaping craving, whereas no such association was observed for block-wise smoking or food craving (see Supplementary Results).

### Classification of High Versus Low Craving

For a brain signature to have clinical relevance or to function as a biomarker, it should be sensitive to both individual difference and cue-dependent variation in normative craving. Hence, we next tested whether the brain signatures could discriminate brain responses to cues associated with the highest versus lowest normative craving levels. We evaluated within-participant forced-choice discrimination by comparing each participant’s responses to cues at the highest versus lowest normative craving levels.

More specifically, we used a forced-choice procedure comparing each participant’s pattern-expression values at their maximum and minimum normative craving levels. All three signatures performed above chance (vaping: 92.4% ± 3.3% SE, binomial test *p* < .001, sensitivity = 92%, specificity = 92%, AUC = 0.95, Figure 1 C; smoking: 98.5% ± 1.5% SE, binomial test *p* < .001, sensitivity = 98%, specificity = 98%, AUC = 1.00, Figure 2 C; food: 93.9% ± 2.9% SE, binomial test *p* < .001, sensitivity = 94%, specificity = 94%, AUC = 0.99, Figure 3 C).

**Figure 2.**
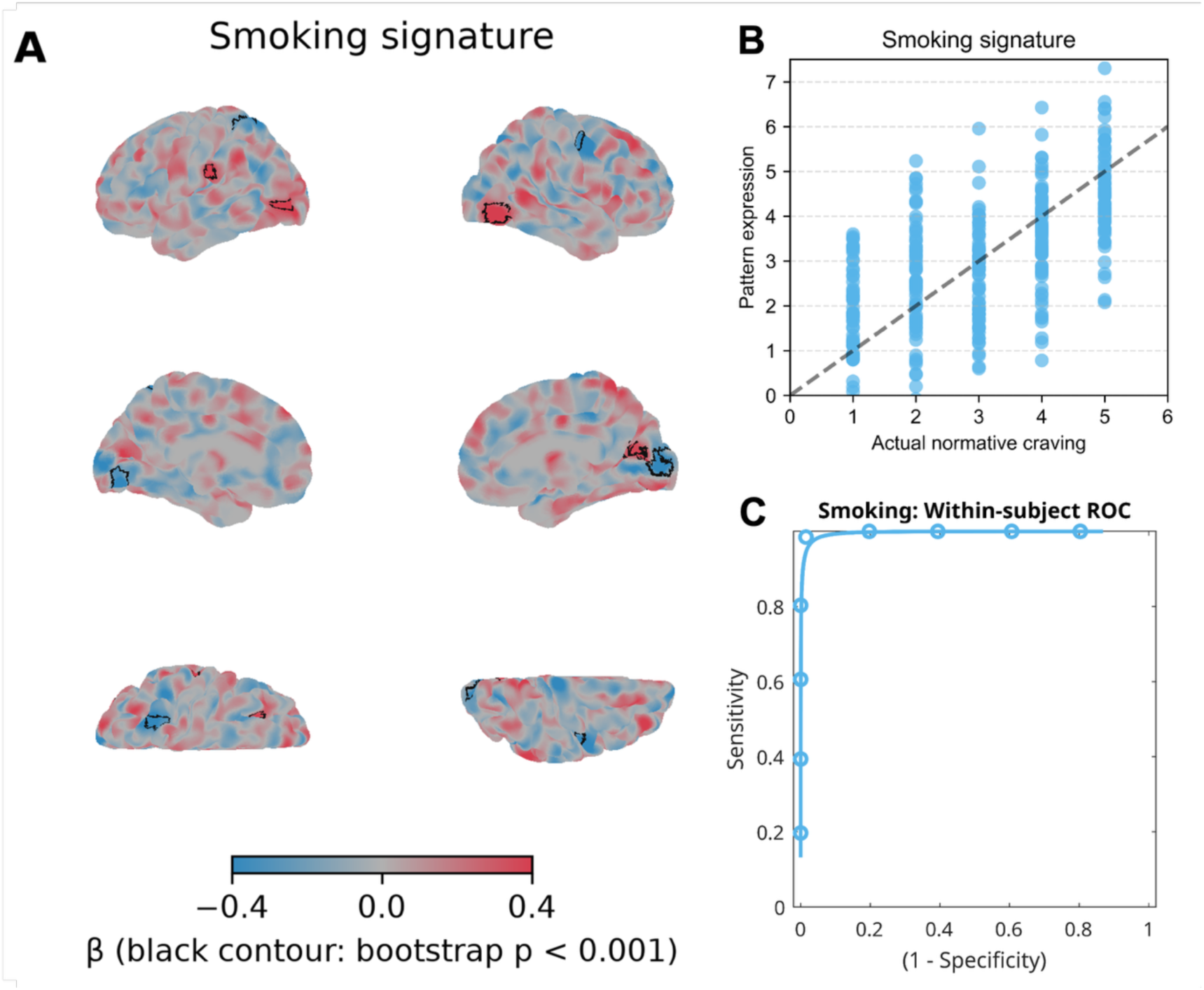
Smoking craving brain signature. **A.** Medial, lateral, and axial views of the brain signature weights. Red and blue indicate positive and negative weights, respectively, with color saturation reflecting the absolute magnitude of the weights. Regions enclosed by black contours survived bootstrap testing at *p* < .001. **B.** Smoking brain signature pattern expression versus normative craving ratings. **C.** Within-subject receiver operating characteristic (ROC) curve for discriminating the highest versus lowest levels of craving.

**Figure 3.**
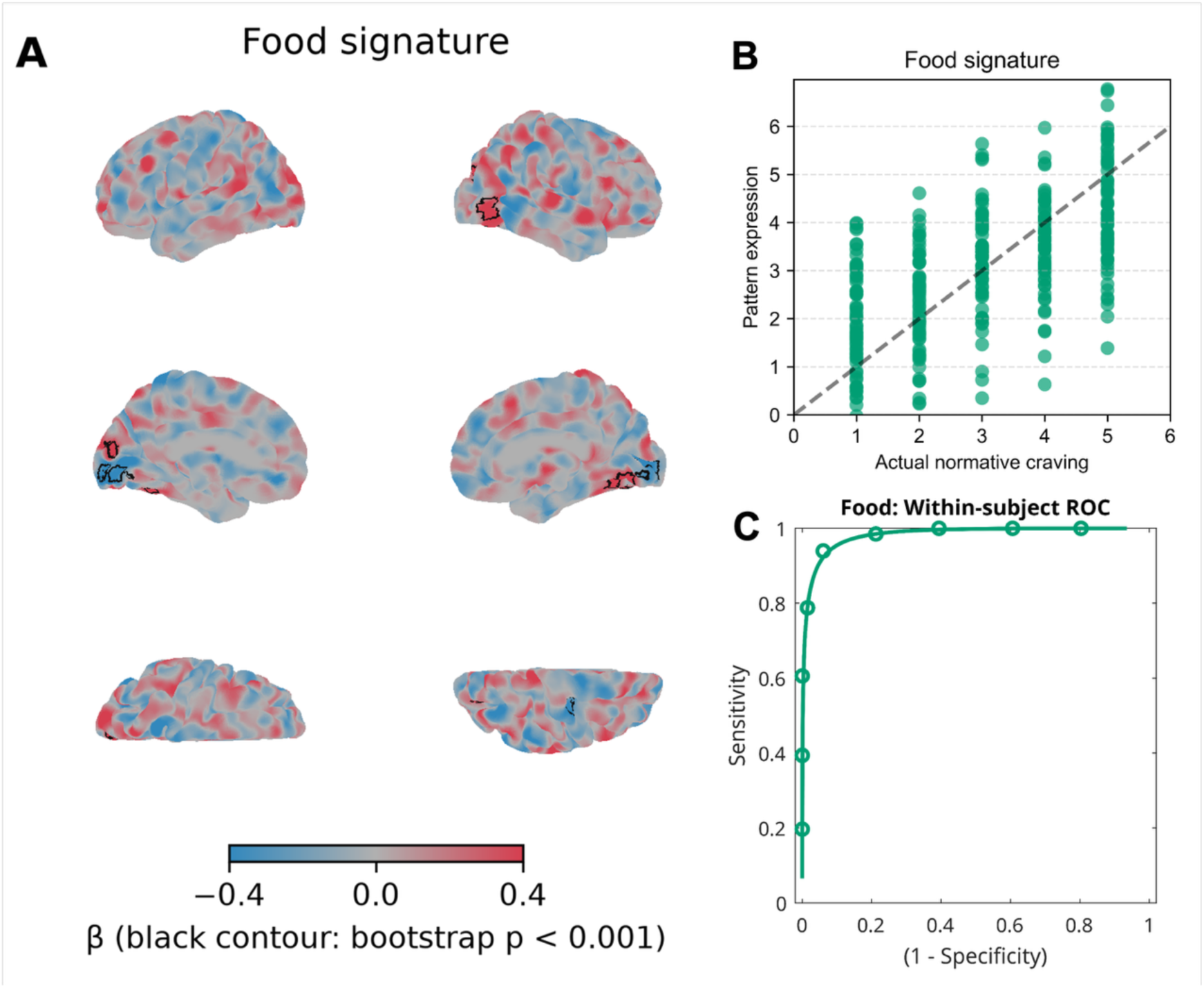
Food craving brain signature. **A.** Medial, lateral, and axial views of the brain signature weights. Red and blue indicate positive and negative weights, respectively, with color saturation reflecting the absolute magnitude of the weights. Regions enclosed by black contours survived bootstrap testing at *p* < .001. **B.** Food brain signature pattern expression versus normative craving ratings. **C.** Within-subject receiver operating characteristic (ROC) curve for discriminating the highest versus lowest levels of craving.

**Figure 4.**
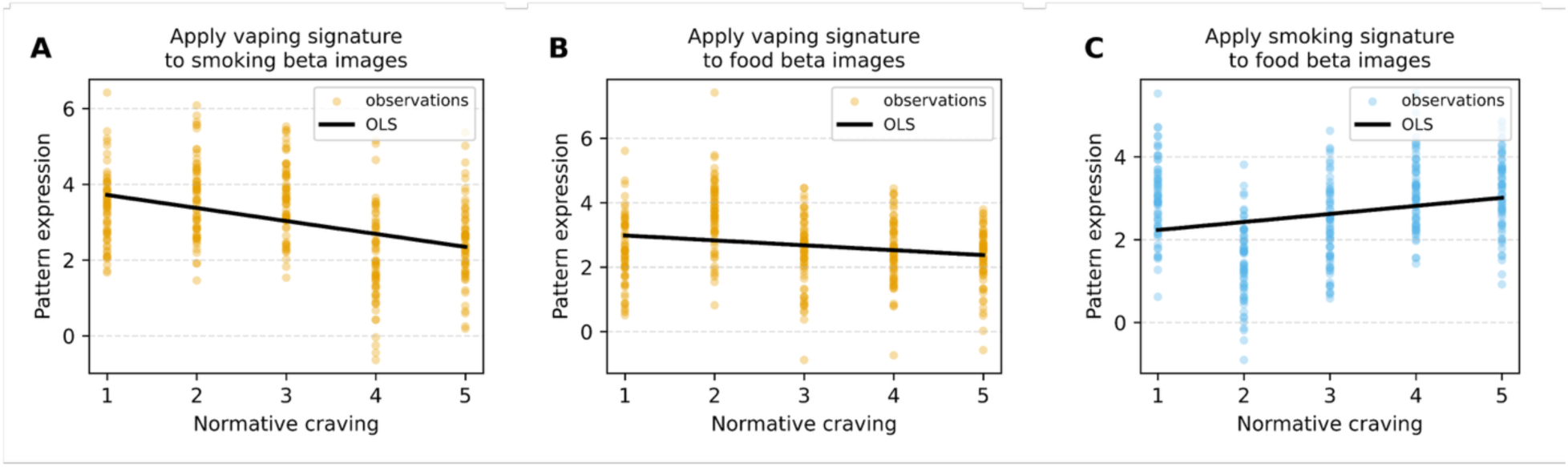
Cross-predictivity tests. **A.** Application of the vaping signature to smoking beta images. **B.** Application of the vaping signature to food beta images. Vaping signature pattern expression decreased as normative craving for smoking and food increased. **C.** Application of the smoking signature to food beta images. Smoking signature pattern expression partially tracked normative food craving. Black lines indicate the ordinary least squares (OLS) linear regression fits.

**Figure 5.**
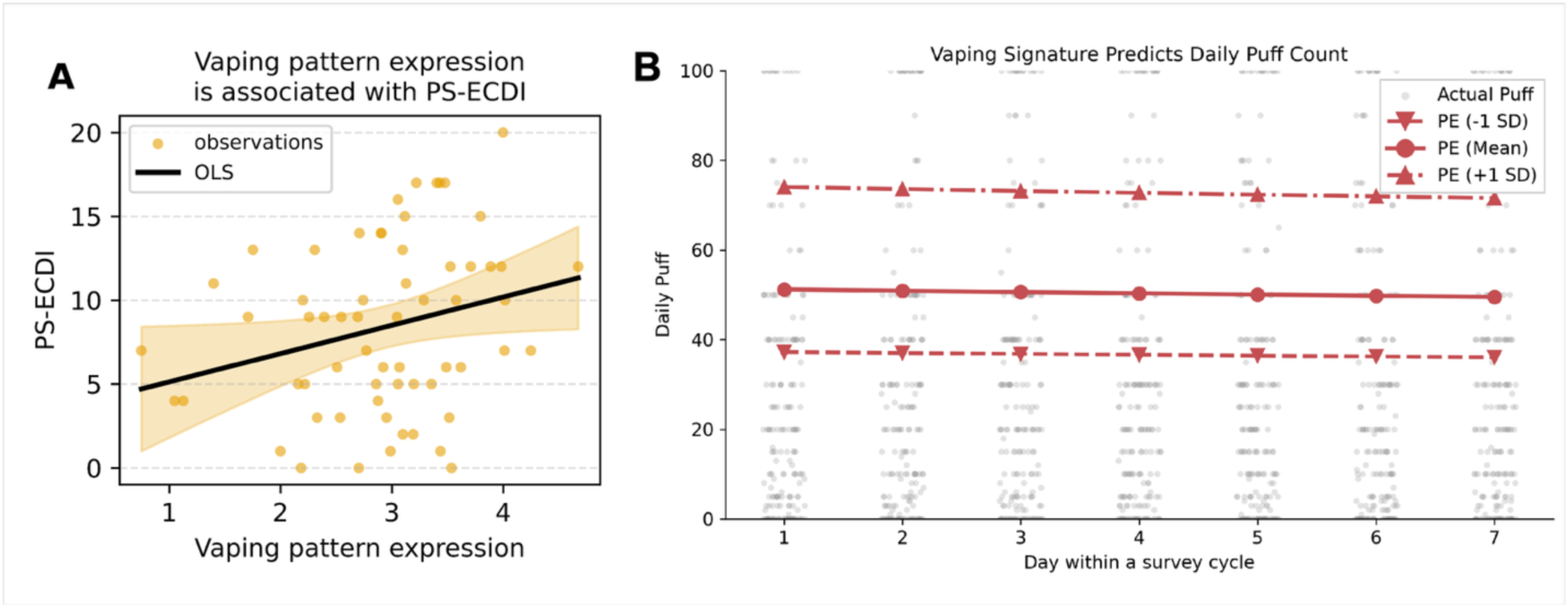
Associations between neural-signature-based predictions and substance use. **A.** Vaping-signature pattern expression was associated with the e-cigarette dependence index, whereas smoking– and food-signature pattern expression showed no significant associations. Black lines indicate the ordinary least squares (OLS) linear regression fits, while yellow dots show the observed values. **B.** Association between vaping-signature pattern expression and daily e-cigarette puff counts. At the end of each of the four weeks following the fMRI scan, participants retrospectively reported their daily e-cigarette use. The x-axis represents the day within each weekly survey cycle, with Day 1 indicating the first day of the week. The line with circles shows model-predicted daily puff counts for participants with vaping-signature pattern expression at the group mean, whereas the lines with upward and downward triangles show predicted puff counts at one standard deviation above and below the group mean, respectively. Observed daily puff counts are shown as gray dots in the background. For visualization purposes, daily puff counts are displayed with an upper cutoff of 100 puffs; the full range of observed values (0–500 puffs) is shown in Figure S1.

**Figure 6.**
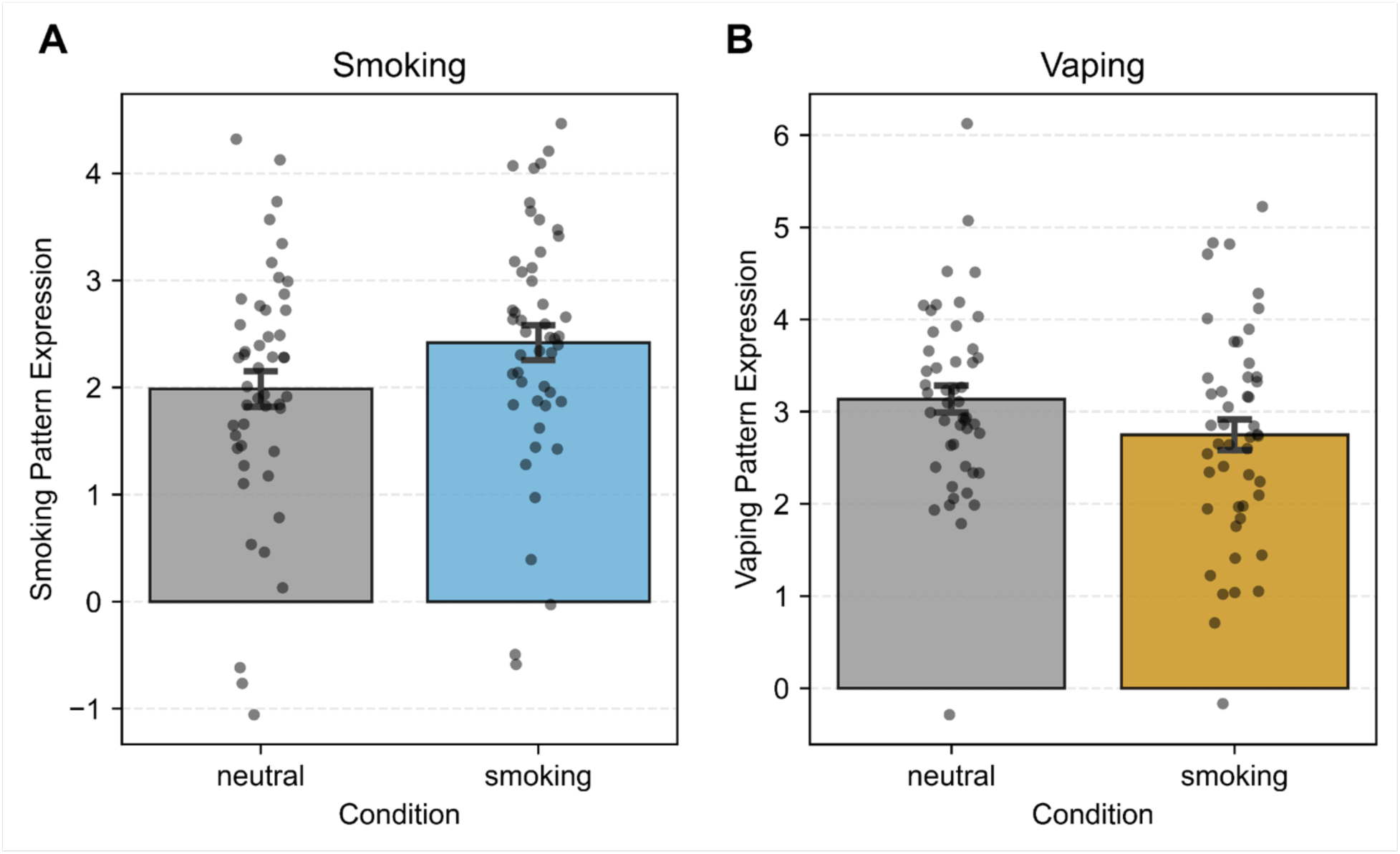
Test nicotine-related signatures on external smoking dataset. **A.** Smoking signature showed higher pattern expression during smoking cue condition, compared to neutral cue condition. **B.** Vaping signature presented the opposite effect, showing higher pattern expression during neutral cue condition vs. smoking cue condition.

### Overlap Between Different Signatures

To what extent do these cue-specific signatures share an underlying neural substrate, and to what extent do they capture unique and distinct neural and psychological constructs? To examine the extent to which the neural craving signatures shared similar distributed spatial patterns, we assessed overlap in two ways: first, by calculating cosine similarity between signatures (see Methods); and second, by applying one signature to predict craving responses elicited by another cue type. Whereas the first approach indexed spatial correspondence, the second indexed practical interchangeability.

We found weak similarity between all pairs of cue-specific signatures: Cosine similarity was 0.03 between the smoking and food signatures, −0.03 between the vaping and smoking signatures, and −0.002 between the vaping and food signatures, where the maximum possible value is 1. Together, these results indicate minimal spatial correspondence among the signatures.

We next tested the overlap between vaping, smoking, and food signatures. For each cross-cue comparison, we applied one signature to the contrast images from the other cue types and regressed normative craving on pattern expression. Results supported limited predictive interchangeability between the vaping signature and the other two signatures, as indicated by negative associations between true smoking and food craving levels and vaping signature pattern expression (smoking: *β* = −0.46 ± 0.06, *t*(328) = −7.83, *p* < .001; food: *β* = −0.24 ± 0.07, *t*(328) = −3.50, *p* = .001). By contrast, smoking and food craving showed partial shared structure, as indicated by a positive association between true food normative craving and smoking signature pattern expression (*β* = 0.30 ± 0.07, *t*(328) = 4.49, *p* < .001).

### Comparison to Other Brain Signatures

How do our cue-specific brain signatures overlap with, or differ from, existing brain signatures? Since brain signatures of food and smoking cue reactivity have been established in prior work (Koban et al., 2023), we first tested whether the food and smoking signatures developed in the present study show correspondence with these previously validated signatures. We then tested whether the vaping signature exhibits a distinct pattern relative to established cue-reactivity signatures. Finally, we examined whether our cue-specific signatures are reducible to brain signatures of related psychological constructs or instead capture cue-specific craving processes.

A closely related contribution is the neural craving signature (NCS) for food and drug craving developed by Koban et al. (2023). In that work, the authors combined neuroimaging data from studies of cigarette, alcohol, and cocaine craving and used machine learning to develop brain signatures that predicted self-reported food and drug craving.

To test whether e-cigarette craving—a substance that is not included in the development of the NCS—could also be captured by the NCS, we applied the signature to the beta image for each cue condition and the control condition. NCS pattern expression was then entered as the dependent variable in a regression model, with cue type included as the predictor. Relative to the control condition, vaping cues elicited significantly greater NCS pattern expression (*β* = 0.40 ± 0.18, *t*(260) = 2.29, *p* = .023), indicating greater predicted craving. Similar effects were observed for smoking cues (*β* = 0.37 ± 0.18, *t*(260) = 2.09, *p* = .037) and food cues (*β* = 0.36 ± 0.18, *t*(260) = 2.04, *p* = .043). These findings indicate that the NCS distinguished vaping, smoking, and food cues from office-supply control cues, suggesting sensitivity to broad cue-related craving rather than specific sensitivity to limited types of drugs that are included in the training set. However, additional analyses indicated that the NCS did not reliably track differences in normative craving intensity within cue categories, particularly during exposure to vaping cues (see the Supplementary Materials for details). A parallel analysis using participants’ block-wise self-reported craving ratings showed a similarly limited and inconsistent pattern of NCS specificity, with only NCS-Food pattern expression tracking block-wise food craving (see the Supplementary Materials for details). Together, these results suggest that the NCS was sensitive to broad, condition-level differences between craving-related and control cues across reward categories but may lack the specificity needed to characterize finer-grained differences in normative craving intensity across vaping cues, highlighting the value of developing a cue-specific brain signature.

Next, to compare our cue-specific signatures with brain signatures of broader psychological constructs, we selected six theoretically relevant signatures, including NCS, and one signature as baseline control. The theoretically relevant signatures captured cognitive control (Kragel et al., 2018; Silvestrini et al., 2020), reward (Speer et al., 2023), negative affect (Chang et al., 2015), and social rejection (Woo et al., 2014). As a control, we included a vicarious pain signature (Krishnan et al., 2016), which captures neural responses to others’ pain. We selected this signature because it reflects secondhand somatic experience but is not directly related to craving.

Our results showed that the vaping signature has low cosine similarity to all existing signatures (vicarious pain cosine similarity = 0.01, other cosine similarities < 0.03, for more information, see Table S3), as did smoking (vicarious pain cosine similarity = –0.02, other cosine similarities < 0.06) and food signatures (vicarious pain cosine similarity = –0.04, other cosine similarities < 0.07). Overall, the present cue-specific weight maps showed little spatial correspondence with the selected signatures of cognitive control, reward, negative affect, social rejection, and vicarious pain. These low similarities do not rule out contributions from these broader processes but suggest that none of the selected signatures closely reproduced the complete spatial patterns identified here.

### Link to Substance Use Behaviors

We next examined whether brain-based prediction of cue-induced craving was associated with e-cigarette dependence severity and subsequent vaping behavior. For each participant, we took the mean leave-one-subject-out pattern-expression value from the corresponding cue-specific signature and defined it as pattern expression. First, we tested the link between pattern expression and participants’ substance dependence severity, assessed using the Penn State Electronic Cigarette Dependence Index (PS-ECDI). Mean-centered age and BMI, as well as sex, were entered in the linear regression model as controls. Sixty-two subjects who had complete data for all covariates were included in the model.

Greater mean vaping-pattern expression was positively associated with e-cigarette dependence severity (vaping: *β* = 1.76 ± 0.86, *t*(57) = 2.05, *p* = .045). In contrast, smoking– and food-signature expression were not significantly associated with e-cigarette dependence (smoking: *β* = −1.28 ± 0.82, *t*(57) = −1.56, *p* = .123; food: *β* = −0.24 ± 1.08, *t*(57) = −0.22, *p* = .828).

We next examined whether expression of the vaping-related brain pattern was associated with real-world e-cigarette use. During the 4 weeks after the fMRI scan, participants reported their daily usage in a series of Timeline Follow-back surveys. A Bayesian Poisson-lognormal mixed-effect model was used to evaluate how standardized vaping pattern expression predicts daily puff counts. Age, BMI, sex, and day within the weekly TLFB cycle were included as covariates. Across 58 participants and 1,407 person-days, greater vaping pattern expression showed a positive but modest prospective association with daily puff rates (*β* = 0.32 ± 0.21; IRR = 1.38, 95% HDI [0.89, 2.05]; posterior *P*[*β* > 0] = .93). On the multiplicative rate scale, a 1-SD increase in vaping pattern expression was therefore associated with an estimated 38% higher expected daily puff rate, although the 95% HDI indicated some uncertainty regarding the precision of the association.

### Proof-of-Principle Cross-Dataset Validation

As a proof-of-principle test of cross-dataset portability, we applied our nicotine-related signatures to an independent archived smoking-cue dataset (Owens et al., 2018, *n* = 48). In that study, participants viewed smoking-related and neutral control cues during fMRI scanning. We applied the vaping and smoking brain signatures to each participant’s smoking and neutral contrast images and computed pattern-expression values for each signature. We hypothesized that the smoking signature would show greater pattern expression for smoking than neutral cues, whereas the vaping signature would not reliably distinguish between the two conditions.

We fit linear mixed-effects models with pattern expression as the outcome, cue condition (smoking vs. neutral), sex, age, and race as fixed effects, and participant as a random intercept. The smoking signature showed the expected effect: pattern expression was higher for smoking than neutral cues (neutral = 1.99, smoking = 2.42; *β* = 0.43 ± 0.09, *z* = 5.02, *p* < .001). The vaping signature showed a significant effect in the opposite direction, with lower pattern expression for smoking than neutral cues (neutral = 3.14, smoking = 2.75; *β* = −0.39 ± 0.15, *z* = −2.56, *p* = .010), indicating non-transfer, and potentially inverse transfer, of the vaping-related predictive pattern to smoking cue reactivity in this sample.

## Discussion

Cue-induced craving is a reliable metric of subsequent substance use, treatment responsiveness and relapse (Nieto et al., 2026, Vafaie & Kober, 2022). However, the application of brain-based inference to e-cigarette craving has been limited by primarily relying on self-reported methods and underpowered brain imaging models. Rather than testing associations between conditions and strength of regional responses on average, we tested whether distributed patterns of brain activity could quantitatively predict craving and whether this representation generalized across individuals, craving states, and datasets.

Our first hypothesis was that distributed brain activity would predict normative craving for e-cigarettes. Consistent with this prediction, the vaping signature showed strong out-of-sample prediction of craving responses to vaping cues. Importantly, the signature did not merely distinguish vaping cues from other types of visually matched craving-inducing cues; rather, its expression tracked variation in craving intensity, demonstrating that the distributed neural activity contains information about shared craving tendency elicited by these cues. This expands previous neuroimaging research on e-cigarette use in an important direction: prior work has primarily characterized vaping cue reactivity by quantifying regional average activation to vaping cues compared to control or other rewarding cues (Chen et al., 2018; Garrison et al., 2018; Grundinger et al., 2025; Liu et al., 2025). Our findings move beyond assessing conditional differences toward out-of-sample prediction: instead of asking where and how the brain responds to vaping cues within a fixed sample, the present approach asks whether patterns of activity across the brain can be used to estimate how much craving those cues evoke in a new observation. This predictive framework provides a reproducible mapping between distributed neural responses and the subjective experience of vaping craving.

The comparison across craving domains further revealed that vaping craving is not simply a subclass of smoking craving or a completely domain-general desire state. Rather, vaping craving appears to be neurobiologically distinct from smoking and food craving, supported by both spatial comparison of the brain signatures and the lack of cross-domain predictive accuracy. Interestingly, high-craving smoking and food cues were associated with lower vaping-signature expression, suggesting that neural expression of vaping craving is attenuated when smoking or appetitive craving is elevated. In contrast, smoking and food craving exhibited comparatively greater similarity in their neural representations and greater cross-predictability, consistent with previous evidence that craving for drugs and foods can recruit substantially overlapping neural systems (Noori et al., 2016; Tang et al., 2012). Together, the representation of vaping craving appears to rely on a meaningfully distinct configuration of distributed neural activity. Further research is needed to characterize the potential suppressive or competitive interactions between vaping-related and other reward-related neural circuits.

Application of the established brain signatures provides an especially informative test of this distinction. As a closely related model, the NCS (Koban et al., 2023) was developed by modeling neural responses to craving for a wide range of substances, as well as palatable food. The NCS successfully distinguished craving-inducing cues (i.e., vaping, smoking, food) from the office-supply control cues in our data, demonstrating that vaping cues engage a broader neural architecture previously associated with craving. However, NCS expression did not reliably predict finer variation in normative craving intensity within the e-cigarette category. This dissociation suggests that a domain-general craving representation may capture whether a motivationally salient craving state is engaged, whereas predicting the intensity of vaping craving requires information encoded in a more domain-specific pattern. Thus, our findings do not imply that vaping craving is neurobiologically unrelated to other forms of craving. Rather, they suggest a nested organization in which vaping engages general craving processes while also recruiting a more specific distributed representation that is necessary for accurately tracking variation in subjective experience.

Regarding our hypothesis of generalizability, the external validation analyses further supported the reliability of brain signatures and their potential applications. When our smoking-craving signature was applied to an independent dataset that recorded smokers’ neural response to smoking and control cues, it successfully distinguished the smoking-cue condition from the control condition, demonstrating that a signature derived from the present modeling framework can transfer across samples and experimental contexts. This provides an important proof of principle for the robustness and portability of craving signatures. At the same time, the vaping-craving signature did not predict amplified craving when smoking cues were presented. The absence of such an effect instead provides additional evidence that the vaping signature captures features of craving specifically associated with e-cigarette use rather than nicotine-related cue reactivity in general. However, a direct external validation of the vaping-craving signature would provide stronger support for this claim. Because research on e-cigarette craving is still at an early stage, future studies should validate the vaping signature using independent vaping cue-reactivity neuroimaging datasets as more such datasets become publicly available.

To explore how brain signatures can advance our understanding of e-cigarette use, we examined associations between brain signature pattern expression and behavioral measures, including self-reported severity of psychological and physical dependence on e-cigarettes and actual use during the 28-day follow-up period. The vaping signature was associated with both measures. Greater pattern expression was associated with greater e-cigarette dependence, indicating that the vaping-craving signature can capture individual differences and may serve as an indicator of e-cigarette-specific nicotine dependence. At a more granular level, exploratory longitudinal analyses of daily vaping behavior provided more tentative evidence in the same direction: greater vaping-signature expression was associated with higher subsequent puff counts. Although the statistical evidence was inconclusive and the estimate was less precise for extreme users, it revealed a notable trend that warrants further investigation. We therefore regard this behavioral association as preliminary. Future studies with larger samples and intensive longitudinal measurements will be important for determining whether neural craving signatures prospectively predict vaping escalation and persistence, and whether these associations differ between moderate users and heavy users.

Several limitations define important directions for future work. First, the signatures were developed using normative craving values reported by an independent sample of vapers, not from the participants who underwent fMRI scans. Although this design strengthens the model’s ability to identify cue-related neural responses that generalize across samples, it inevitably reduces its sensitivity to individual-level idiosyncrasies. Future work could adopt hierarchical approaches that estimate shared neural representations while permitting person-specific deviations.

Second, we propose that future studies should test brain signatures on various forms of cues and utilize additional craving measures to establish convergent validity. The current paradigm relied primarily on visual vaping cues. Visual images offer strong experimental control and enable direct comparisons across cue categories, but real-world craving is elicited by a richer combination of sensory and contextual information. Moreover, normative craving served as the primary criterion for developing brain signatures in this study. Although self-report remains indispensable when studying an experiential state, it can be influenced by introspective ability, habituation, social desirability, and deception (Caputo, 2019). Future studies can strengthen brain signatures’ external validity by incorporating different modules of cues, including videos, personalized cues and actual vaping opportunities, as well as calibrate brain signatures using physiological responses, biomarkers of nicotine exposure, and ecological momentary assessments.

Finally, the clinical validity of the vaping signature remains to be established. A brain signature becomes particularly valuable as a biomarker when it can predict outcomes that matter beyond the scanning session. Building on our exploratory findings that link brain signature expression to behavior, future longitudinal studies should test whether vaping-signature expression predicts subsequent clinical trajectories, including escalation of use, development of nicotine dependence, quit attempts, relapse, or transition to combustible cigarette use.

In conclusion, the present study provides three complementary advances in understanding e-cigarette craving. First, distributed patterns of brain activity reliably predicted the intensity of vaping craving. Second, vaping craving exhibited a neural representation that was substantially distinct from smoking and food craving and could not be fully captured by an established domain-general craving signature. Third, the successful prediction of smoking craving in an independent dataset demonstrates that neural craving signatures developed using this pipeline can generalize beyond the original sample. Together, these findings establish a reproducible neural model of vaping craving and provide a foundation for linking the neural activity of e-cigarette craving to nicotine dependence, real-world use, and ultimately, clinically relevant outcomes.

## Method

### Participants

#### fMRI sample

The fMRI sample included 66 nonsmoking young adult vapers aged 18–29 years (*M* = 20.08, SD = 1.50; 46 women, 20 men). Participants self-identified as European American (*n* = 49), African American (*n* = 3), Asian (*n* = 9), or multiracial/other race (*n* = 5); eight participants identified as Hispanic. Eligibility required vaping on at least 15 of the past 30 days and no current cigarette smoking, defined as responding “0” to the question, “During the past 30 days, on how many days did you smoke cigarettes, even one or two puffs?” Exclusion criteria included a history of major medical, psychiatric, or neurological conditions, and any contraindication for research MRI, including certain metal implants, pregnancy, or claustrophobia. All participants provided written informed consent. The study was approved and monitored by the University of Georgia Institutional Review Board.

#### Image annotation sample

An independent sample of 442 participants was recruited for the image annotation task from CloudResearch Connect, an online participant recruitment platform. Participants were current regular vapers, defined as vaping at least weekly during the past 30 days. The sample was recruited using quotas designed to match the U.S. young adult population on gender, race, and ethnicity, based on U.S. Census data. Participants were 18–29 years old (*M* = 24.69, SD = 3.24). Of the analytic sample, 214 (48.42%) identified as women, 104 (23.53%) identified as Hispanic, 73.30% identified as European American, 14.93% identified as African American, and 11.76% identified as another race or ethnicity. Participants were excluded if they did not complete the full study, including the questionnaire battery and rating tasks; failed an attention check; failed the CAPTCHA bot check; or showed evidence of straightlining. Participants who met any exclusion criterion were excluded from the final analyses. All participants provided informed consent, and the annotation study was approved by the University of California Santa Barbara Institutional Review Board.

### Stimuli

The stimulus set consisted of 144 images across four categories: smoking-related, vaping-related, food-related, and office-supply images, with 36 images per category. Office-supply images served as control stimuli. Images were designed to minimize differences in low-level and compositional visual properties across categories, including color, size, layout, and object position (e.g., a single e-cigarette was paired with a single stapler). A series of control analyses (Supplementary Results; Figures S2 and S3) showed that cue categories differed primarily in color, whereas normative craving levels were largely unconfounded with low-level visual features. Presentation format was matched as well, including whether objects appeared alone, in groups, or in interaction with people. Gender representation and the presence of vapor, smoke, or steam were systematically matched across corresponding images in experimental conditions.

### Image annotation task

The image annotation task was preregistered [public link] and was designed to obtain normative craving ratings for all stimuli used in the fMRI task. Each participant viewed a subset of images drawn from the full stimulus pool. Image sets included a balanced number of smoking-related, vaping-related, food-related, and control images. On each trial, one image was presented on the screen, and participants rated their immediate craving in response to the image using three items: “After seeing this image, how much do you want to vape/smoke/eat right now?” Responses were made on a 0–100 scale, with 0 indicating not at all and 100 indicating extremely. To reduce the influence of extreme values and harmonize the rating scales, ratings for each craving dimension—vaping, smoking, and eating—were discretized into five levels based on the 20th, 40th, 60th, and 80th percentiles of the corresponding rating distribution. These craving-level labels were used in subsequent analyses. Descriptive statistics of normative craving ratings can be found in Supplementary Table 1 and 2.

### Procedure of fMRI study

Before the fMRI session, participants completed a survey assessing sociodemographic characteristics and vaping-specific nicotine dependence using the Penn State Electronic Cigarette Dependence Index (PS-ECDI) (Foulds et al., 2015), a 10-item measure of e-cigarette dependence.

The 1-hr fMRI session included two imaging runs of a visual cue-reactivity task. Each run lasted 414 s and included: three 30-s blocks for each cue condition (vaping, smoking, and food), three 20-s blocks of control images (office supplies), and three 20-s blocks of scrambled images. Block order was pseudorandomized. Within each cue block, six distinct images were presented for 5 s each.

Between blocks, participants rated their craving for vaping-, smoking-, and food-related products on a 7-point Likert scale ranging from 1 (not at all) to 7 (very much). There were four rating sessions during each run. Specifically, participants rated e-cigarette craving after the first block of control images and the first vaping-cue block, cigarette craving after the first smoking-cue block, and food craving after the first food-cue block. These ratings were originally included to maintain participant engagement and were additionally used in supplementary analyses examining associations between neural-signature expression and participants’ self-reported block-wise craving.

After the fMRI scan, participants self-reported their daily e-cigarette use via follow-up surveys administered every seven days for 4 weeks, following the Timeline Follow-Back (TLFB) approach (L. Sobell & M. Sobell, 1996; Brown et al., 1998; Griffith et al., 2009). TLFB is a calendar-based retrospective assessment designed to facilitate recall of daily substance use. In each follow-up survey, participants were presented with each of the preceding seven days and asked to report the number of e-cigarette puffs they had taken on each day. Calendar dates and temporal prompts were provided to help participants reconstruct their daily use by anchoring recall to specific days and events. Sixty-two participants completed at least one week’s survey, while forty-one of them completed all surveys.

### fMRI Data Acquisition and Preprocessing

fMRI data were collected at the University of Georgia using a GE Discovery 750 3T MRI scanner with a 32-channel head coil (GE Healthcare, Milwaukee, WI). Blood oxygenation level– dependent (BOLD) fMRI data were acquired using a whole-brain, single-shot gradient-echo echo-planar imaging sequence with the following parameters: repetition time (TR) = 2,000 ms; echo time (TE) = 25 ms; field of view (FOV) = 225 × 225 mm²; matrix = 64 × 64; slice thickness = 3.5 mm; flip angle = 90°; 40 axial slices; and effective voxel resolution = 3.52 × 3.52 × 3.50 mm³. Following BOLD fMRI acquisition, structural images were acquired using a 3D fast spoiled gradient-echo sequence with the following parameters: TR = 7.80 ms; TE = 2.98 ms; FOV = 256 × 256 mm²; matrix = 256 × 256; 164 sagittal slices; flip angle = 20°; and effective voxel resolution = 1 × 1 × 1 mm.

fMRI data were preprocessed and analyzed using SPM12 (Wellcome Centre for Human Neuroimaging, 2014) and the ArtRepair toolbox (Mazaika et al., 2005) in MATLAB (MathWorks, Natick, MA). Functional images underwent slice-wise outlier detection and repair using art_slice, slice-timing correction, realignment and head-motion correction, coregistration of structural and functional images, normalization to Montreal Neurological Institute (MNI) space with 3-mm isotropic voxels via unified segmentation, spatial smoothing with an 8-mm full-width at half-maximum Gaussian kernel, and volume-wise outlier correction using art_global.

### fMRI Data Analysis

For each participant, separate first-level GLMs were estimated for each cue category. For vaping, smoking, and food cues, cue events were modeled across two runs with separate regressors corresponding to five normative craving levels derived from the independent image annotation task. Event durations were modeled according to the task design. Three additional regressors modeled variance associated with non-target cues, scrambled images, and rating periods. Six rigid-body motion parameters were included as nuisance regressors. A 128-s high-pass filter and AR(1) autocorrelation correction were applied. This procedure yielded five contrast images of interest per run for each cue category and participant, with one contrast image corresponding to each craving level.

### Training and Cross-Validation of Brain Signatures

The contrast images corresponding to the five craving levels were used to train a linear predictive model of craving using least absolute shrinkage and selection operator–principal component regression (LASSO-PCR) (Tibshirani, 1996), implemented with the CANlab toolbox (Cognitive and Affective Neuroscience Laboratory, 2024). LASSO-PCR is well suited for predicting continuous outcomes from high-dimensional neuroimaging data, where the number of potential predictors (i.e., voxels) substantially exceeds the number of observations and where predictors are often highly correlated across brain regions and functional networks (Chang et al., 2015; Wager et al., 2013; Zhou et al., 2021). The approach reduces overfitting by first applying principal component regression to summarize covarying voxel patterns into lower-dimensional components. It then applies LASSO regularization, which shrinks the weights of less stable or less predictive components toward zero. Voxel-level weights can subsequently be reconstructed from the component weights, yielding an interpretable whole-brain predictive map that can be applied to external datasets.

For each cue type, we trained and evaluated a LASSO-PCR model using leave-one-subject-out cross-validation. In each fold, all contrast images from one participant, corresponding to the five craving levels, were held out together to ensure independence between the training and test sets at the participant level. The model was trained on data from the remaining participants and then applied to each held-out contrast images. Pattern expression was computed as the inner product between the trained weight map and the held-out contrast image, plus the scalar intercept estimated during training. This procedure yielded an out-of-sample pattern-expression score for each craving level and participant. Model performance was evaluated in Multiple Linear Regression models, which control for sex, age, BMI, and average head motion.. Because each participant contributed repeated observations across five craving levels, we estimated cluster-robust standard errors at the participant level to account for within-subject dependence. This approach preserves the coefficient estimates while adjusting standard errors, confidence intervals, and statistical inference for potential dependency among observations from the same participant. Regressions with cluster robust variance estimators are potentially advantageous compared to linear mixed-effect models, since they rely on weaker assumptions (Lee & Pustejovsky, 2024), and has been widely applied in empirical studies that involve repeated behavioral observations (Phillips et al., 2025; Orchinik et al., 2025; Levine et al., 2018). Separate models were trained for vaping, smoking, and food cues using only the contrast images from the corresponding cue category.

### Bootstrapping and Thresholding

To assess the reliability of voxel-level weights, we performed a bootstrap analysis on the full sample using the CANlab predict function with cv_lassopcr, five-fold cross-validation, and 10,000 bootstrap resamples (bootsamples). For each voxel, bootstrap Z scores were computed as the mean weight across resamples divided by the standard deviation across resamples; two-tailed *p* values were obtained from the standard normal distribution. For visualization, uncorrected *p* < .001 (minimum cluster size = 10 voxels) threshold maps are overlaid on top of threshold maps. Unthresholded maps were used for all prediction analyses.

### Classification Analyses

We used receiver operating characteristic (ROC) analyses to evaluate whether cue-specific brain signatures discriminated high versus low craving using leave-one-subject-out cross-validated pattern-expression values, implemented with the CANlab ‘roc_plot’ function. Area under the curve (AUC) was reported as a threshold-independent measure of discrimination, and binomial tests were used to assess the statistical significance of classification accuracy.

The same brain signature’s cross-validated pattern-expression values were evaluated under a within-participant classification scheme, distinguishing cues assigned to the highest (level-5) versus lowest (level-1) normative craving levels. High-craving observations were treated as the positive class and low-craving observations as the negative class.

We compared each participant’s pattern-expression values for cues at the highest and lowest normative craving levels in a two-alternative forced-choice procedure. For each participant, we computed the paired difference d_s as the high-craving pattern-expression value minus the low-craving pattern-expression value. The two scores were then expressed relative to their pair mean, yielding centered values of d_s/2 for the high-craving observation and −d_s/2 for the low-craving observation. The forced-choice decision used a fixed threshold of zero, so a participant was classified correctly when the high-craving score exceeded the low-craving score. Reported sensitivity was calculated as the proportion of high-craving centered scores at or above zero, and specificity as the proportion of low-craving centered scores below zero. Because centering makes these two classifications complementary around the same zero threshold, sensitivity and specificity are numerically equivalent in this analysis. For the ROC curve, candidate thresholds were swept across the range of these centered scores to estimate threshold-dependent true– and false-positive rates and AUC.

### External Dataset

The external smoking dataset (Owens et al., 2018) included 51 participants. One participant was missing one scanning run, and two participants were excluded because of excessive head motion, defined as framewise displacement greater than 0.35 mm in more than 33% of brain volume acquisitions. The final analytic sample included 48 participants. For more details about this dataset, see Supplementary Materials.

### Application of Brain Signatures to the External Dataset

A first-level GLM was estimated in SPM12 across the two functional runs. Smoking and neutral cues were modeled as separate regressors within each run, and an additional regressor modeled the rating periods. Rest periods served as the baseline. To align with the original analysis by Owens et al. (2018), 18 motion parameters were included as nuisance regressors. These included the six rigid-body realignment parameters expanded to linear, quadratic, and cubic terms. Spike regressors were also included for motion-censored volumes, and participants with more than 33% of volumes censored were excluded from analysis. A 128-s high-pass filter and AR(1) autocorrelation correction were applied. Two contrast images were extracted for each participant, representing neural responses to smoking and neutral cues, respectively.

After extracting contrast images for smoking and neutral cues, we applied the brain signatures developed in the primary dataset to the external dataset. Each contrast image was normalized using L2 normalization to exclude scaling effects and then projected onto the brain signature. Pattern expression was computed as the dot product between the normalized contrast image and the brain signature weight map, plus the scalar intercept estimated during model training. This procedure was conducted separately for each participant and cue type, yielding one pattern-expression value for smoking cues and one pattern-expression value for neutral cues per participant.

### Cosine Similarity Between Brain Signatures

To quantify the spatial correspondence between brain signatures, we computed cosine similarity between their voxel-wise weight vectors. For each pair of signatures, analysis was restricted to voxels with nonzero weights in both maps, ensuring that similarity reflected shared spatial support rather than sparsity mismatch. Cosine similarity was defined as the dot product of the two weight vectors divided by the product of their L2 norms, yielding values from −1 (opposite directions) to 1 (identical directions), with 0 indicating orthogonality.

### Bayesian Poisson-Lognormal Mixed-Effect Model of Daily Puff Counts

To test whether vaping-related neural pattern expression was associated with real-world e-cigarette use, we analyzed daily puff counts reported during the 28-day Timeline Follow-Back (TLFB). For each participant, vaping pattern expression was defined as the mean leave-one-out predicted vaping craving score and standardized across participants, while age, BMI, sex, and day within the weekly TLFB cycle were included as covariates. The model included a participant-specific random intercept. Age and BMI were centered at sample means, and sex was coded with female as the reference category. Analyses were restricted to observations with non-missing puff count, pattern expression, age, BMI, and sex, which yielded a sample of 58 participants. For further details about model specification, see Supplementary.

