## Supplementary Materials for "A brain signature of vaping cue-elicited craving"

#### Supplementary Results

**Descriptive statistics of normative craving ratings**

| Cue | Mean | SD | Min | Max |
|---|---|---|---|---|
| Vaping | 55.72 | 6.82 | 39.23 | 68.93 |
| Smoking | 41.35 | 6.45 | 28.00 | 58.50 |
| Food | 59.38 | 6.80 | 46.53 | 72.07 |

Table S1. Descriptive statistics of normative craving ratings (0-100 scale) for each cue category, computed across the 36 stimuli per condition used in the cue-reactivity task.

| Cue | Level 1 | Level 2 | Level 3 | Level 4 | Level 5 |
|---|---|---|---|---|---|
| Vaping | 39.23–50.82 | 51.05–54.00 | 54.06–57.75 | 57.90–60.76 | 61.02–68.93 |
| Smoking | 28.00–35.80 | 35.88–40.37 | 40.61–43.37 | 43.45–46.03 | 47.06–58.50 |
| Food | 46.53–53.21 | 53.46–57.64 | 57.71–59.94 | 60.50–67.03 | 67.12–72.07 |

Table S2. Range of raw normative craving rating scores corresponding to each of the five quantile-based craving levels, binned separately within each cue condition.

**Brain signature performance: model comparison**

We compared three models to evaluate the brain signatures' predictive performance: (A) a multiple linear regression model in which normative craving was the dependent variable and the brain signature pattern expression value was the independent variable, controlling for the effects of sex, age, body mass index (BMI), and average head motion, with cluster-robust standard errors clustered by participant to account for each participant contributing five observations (62 clusters; t-tests with df = 61); (B) a linear mixed-effects model with the same fixed effects plus a random intercept for subject; and (C) a linear mixed-effects model with the same fixed effects, a random intercept for subject, and an additional random slope for pattern expression by subject.

The strong positive association between pattern expression and normative craving was stable across all three models and across all three cue conditions (vaping, food, smoking): $\beta$ ranged from 0.52 to 0.66 (all $p < .001$) and changed negligibly whether or not random effects were included. By contrast, neither the subject random intercept in Model B nor the random slope in Model C explained meaningful variance: the estimated random-intercept variance was approximately 0 in every condition, likely due to the shared normative craving levels. The random-slope variance and intercept–slope correlation were likewise negligible or degenerate (correlation at the ±1 boundary), indicating singular fits.

Consistent with this, the two more complex models, B and C, did not outperform the multiple linear regression model: BIC was lowest (best) for Model A in all three conditions (vaping: 1038.7 vs. 1044.4 vs. 1055.0; food: 973.0 vs. 978.7 vs. 990.2; smoking: 980.1 vs. 985.8 vs. 997.3, for Models A, B, and C respectively), reflecting the added parameter penalty of the random effects without a corresponding gain in model likelihood. We therefore report results from the simpler multiple linear regression model (Model A) in the main text.

**Cosine similarity between signatures**

| Signature | Vaping | Smoking | Food |
|---|---|---|---|
| VPS | 0.009 | -0.015 | -0.042 |
| Kragel18CogControl | 0.011 | 0.043 | 0.003 |
| Stroop | 0.023 | -0.052 | 0.062 |
| Rejection | 0.019 | 0.033 | 0.006 |
| PINES | -0.005 | 0.057 | 0.011 |
| NCS | -0.028 | -0.015 | -0.008 |
| BRS | 0.007 | 0.031 | 0.002 |

Table D3. Values in the cells show the cosine similarity between vaping, smoking and food signature with established signatures. VPS: Vicarious Pain Signature (Krishnan et al., 2016); Kragel18CogControl: Cognitive Control signature (Kragel et al., 2018); Stroop: Stroop-demand SVM pattern (Silvestrini et al., 2020); Rejection: Romantic Rejection signature (Woo et al., 2014); PINES: Picture-Induced Negative Emotion Signature (Chang et al., 2015); NCS: Neural Craving Signature (Koban et al., 2023); BRS: Brain Reward Signature (Speer et al., 2023).

**Test vaping brain signature against block-wise ratings**

To further assess whether the vaping signature tracks participants' self-reported block-wise craving, we applied the signature to block-level beta images obtained during vaping-, smoking-, and food-cue blocks. For each cue block, we calculated the expression of the vaping signature and entered these values into a linear regression model predicting the craving rating reported immediately after the block. Because craving ratings were collected only after the cue blocks were first presented, this analysis was necessarily restricted to three blocks, with each block-level beta image paired with a single rating reflecting the participant's craving at the end of that block. This design therefore assessed craving at a coarser temporal resolution than the stimulus-level analysis and, by relying only on the rated blocks, did not capture signature–craving associations across the full one-hour scanning session.

We built simple linear regression models using each signature's pattern expression to predict block-wise cravings, with standard errors clustered by participant. Vaping-signature expression showed a positive association with participants' block-wise vaping craving, although the association did not reach conventional statistical significance ($\beta = 0.21 \pm 0.11$, $t(59) = 1.90$, $p = .062$). The magnitude of this association was smaller than that observed in the primary normative-craving analysis, possibly due to different sampling resolutions. In contrast, vaping-signature expression did not significantly predict craving during smoking-cue or food-cue blocks (smoking: $\beta = 0.00 \pm 0.05$, $t(55) = 0.01$, $p = .994$; food: $\beta = -0.09 \pm 0.10$, $t(58) = -0.87$, $p = .390$). Together, these exploratory findings provide tentative evidence that vaping-signature expression covaries with participants' self-reported vaping craving, whereas no significant associations were observed for smoking- or food-cue craving.

**Test NCS against block-wise rating**

To examine whether established NCS models track participants' self-reported block-wise craving craving, we applied the general NCS, NCS-Drug, and NCS-Food signatures (Koban et al., 2023) to our individual GLM beta images. Following Cortes et al. (2026), we computed block-level pattern expression and regressed block-level self-reported craving ratings on pattern expression. Age, sex, BMI, and mean head motion are used as control variables; standard errors were clustered by participant.

NCS-Food pattern expression was positively associated with participants' block-wise food craving ratings in our study ($\beta = 0.29 \pm 0.13$, $t(55) = 2.26$, $p = .028$) but was not significantly associated with block-wise vaping- or smoking-craving ratings (vaping: $\beta = -0.08 \pm 0.15$, $t(56) = -0.56$, $p = .581$; smoking: $\beta = -0.02 \pm 0.09$, $t(53) = -0.25$, $p = .805$), suggesting limited cross-domain correspondence in this sample. General NCS predictions were not significantly associated with self-reported craving for any cue type (all $p$s > .23). NCS-Drug pattern expression was not significantly associated with smoking- or food-craving ratings (both $p$s > .47). Moreover, its predictions for vaping craving were negatively associated with true vaping craving ($\beta = -0.51 \pm 0.20$, $t(56) = -2.53$, $p = .014$), meaning that higher self-reported craving corresponded to lower NCS-Drug–based predictions.

**Test NCS against normative rating**

Next, we examined whether established NCS models track normative craving by applying the general NCS, NCS-Drug, and NCS-Food signatures to beta images corresponding to normative vaping, smoking and food craving. We regressed cue-specific craving ratings on NCS, NCS-Drug and NCS-Food pattern expression, controlling for age, sex, BMI, and mean head motion, with standard errors clustered by participant (62 clusters; df = 61).

NCS-Drug pattern expression was positively associated with smoking normative rating ($\beta = 0.20 \pm 0.06$, $t(61) = 3.48$, $p < .001$), but this effect was absent when comparing NCS-Drug expression to vaping- and food- normative rating (vaping: $\beta = -0.24 \pm 0.08$, $t(61) = -3.19$, $p = .002$; food: $\beta = 0.02 \pm 0.07$, $t(61) = 0.30$, $p = .764$), suggesting that the distributed brain pattern captured by NCS-Drug tracks smoking craving selectively. General NCS was not positively significantly associated with normative craving for any cue type (vaping: $\beta = -0.11 \pm 0.05$, $t(61) = -2.30$, $p = .025$; smoking: $\beta = 0.01 \pm 0.05$, $t(61) = 0.26$, $p = .793$; food: $\beta = -0.07 \pm 0.04$, $t(61) = -1.55$, $p = .127$). NCS-Food pattern expression was positively associated only with smoking normative rating ($\beta = 0.11 \pm 0.05$, $t(61) = 2.25$, $p = .028$), but not with food or vaping normative rating (food: $\beta = -0.05 \pm 0.05$, $t(61) = -0.95$, $p = .346$; vaping: $\beta = -0.15 \pm 0.06$, $t(61) = -2.72$, $p = .009$).

**Low-level visual features of cue stimuli**

Of the 17 low-level visual features, eight were retained after the two-step redundancy pruning described in the Supplementary Method (final VIFs 1.51–3.28): mean luminance, RMS contrast, CIELAB a*, CIELAB b*, colorfulness, entropy, GLCM correlation, and spectral slope. Images sharing a stimulus number were substantially dependent (median ICC = 0.42, range 0.11–0.77), demonstrating a successful matching procedure during stimuli development. To account for the stimulus number dependency, we used paired comparisons across stimulus numbers for the category post hoc tests.

Cue categories differed mainly in color (Figure S2). Category effects were significant after FDR correction for CIELAB a* ($F(2, 93) = 7.17$, $q = .003$, $\eta_p^2 = .13$), CIELAB b* ($F(2, 93) = 11.63$, $q < .001$, $\eta_p^2 = .20$), and colorfulness ($F(2, 93) = 11.78$, $q < .001$, $\eta_p^2 = .20$). Paired comparisons showed that food images were redder (higher a*; vs. vaping: $d_z = 0.56$; vs. smoking: $d_z = 0.54$; both $p_{Holm} = .006$) and yellower (higher b*; vs. vaping: $d_z = 0.88$; vs. smoking: $d_z = 0.71$; both $p_{Holm} < .001$) than vaping and smoking images, and colorfulness increased from vaping to smoking to food images (all pairwise $p_{Holm} \leq .004$). GLCM correlation, Luminance, RMS contrast, and entropy did not differ across categories (all $q > .40$).

Craving level had no significant main effect on any feature (Figure S3; all $q > .29$), and no feature showed a significant overall linear trend across craving levels (all $q > .53$). The only craving-related effect involved RMS contrast: the category × craving level interaction did not survive FDR correction ($F(8, 93) = 2.80$, $p = .008$, $q = .065$), but the linear trend across craving levels differed between categories ($F(2, 102) = 7.82$, $q = .006$). RMS contrast increased with craving level among smoking images ($b = 0.42$ SD per level, $q = .009$) but not among vaping ($b = -0.18$, $q = .68$) or food images ($b = 0.05$, $q = .94$). Together, cue categories differed primarily in color, whereas normative craving levels were largely unconfounded with low-level visual features.

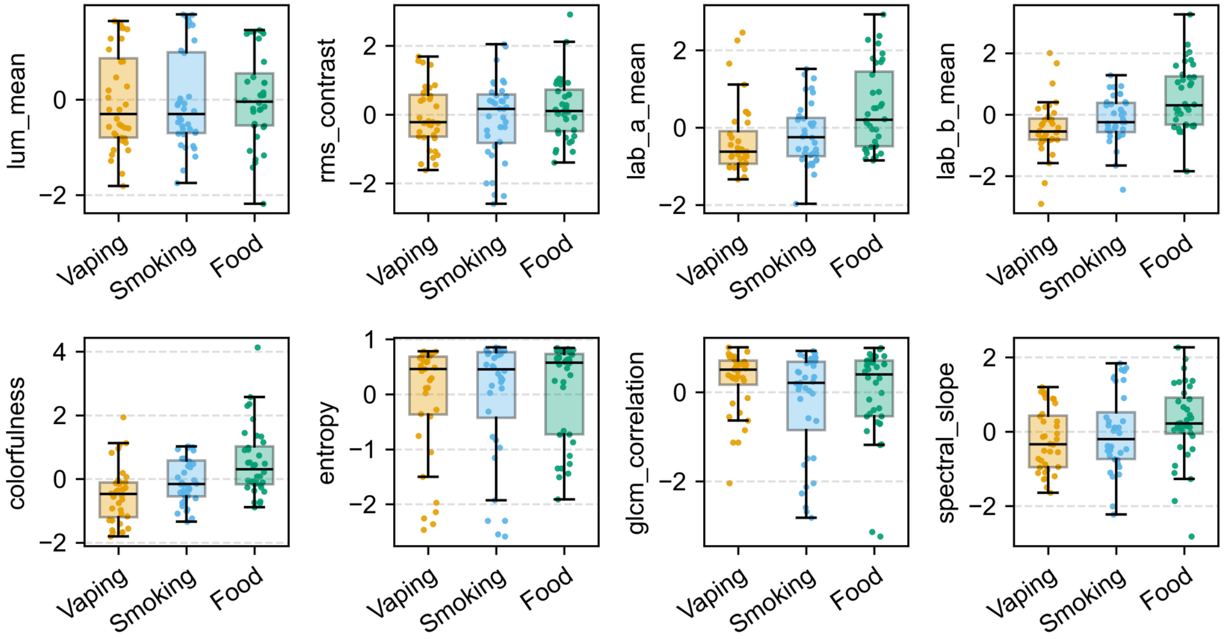


**Figure S2. Low-level visual features of the cue images by category.** Each panel shows one of the eight features retained after redundancy pruning, z-scored across the 108 cue images. Boxes show the median and interquartile range, whiskers extend to 1.5 × the interquartile range, and points show individual images (n = 36 per category). Panel labels: lum_mean, mean luminance (CIELAB L*); rms_contrast, root-mean-square contrast; lab_a_mean, mean CIELAB a* (green–red axis); lab_b_mean, mean CIELAB b* (blue–yellow axis); colorfulness, colorfulness index (Hasler & Süsstrunk, 2003); entropy, Shannon entropy of the grayscale histogram; glcm_correlation, gray-level co-occurrence matrix (GLCM) correlation; spectral_slope, slope of the radially averaged amplitude spectrum in log–log coordinates.

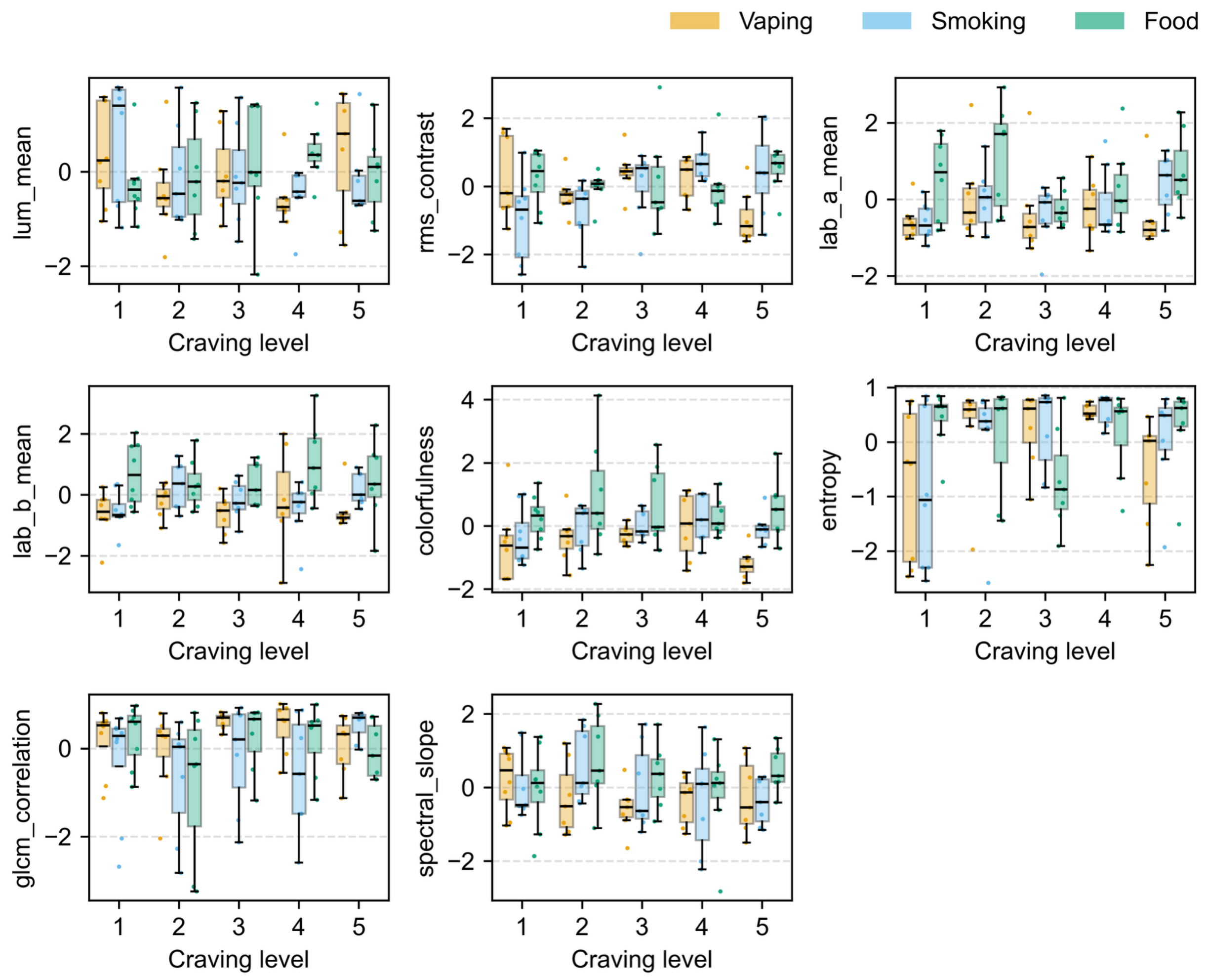


**Figure S3. Low-level visual features of the cue images by craving level.** Each panel shows one of the eight retained features (z-scored across the 108 cue images; panel labels as in Figure S2) across the five normative craving levels, which were binned within category. Within each craving level, boxes are shown separately for vaping (orange), smoking (blue), and food (green) images. Boxes show the median and interquartile range, whiskers extend to 1.5 × the interquartile range, and points show individual images ($n$ = 7–8 per category and craving level). Visual feature labels are the same as Figure S2.

### Supplementary Method

##### Bayesian Poisson-lognormal mixed-effect model

We defined a Bayesian Poisson-lognormal mixed-effect model to test the association between vaping signature pattern expression and daily puff counts reported during the 28-day Timeline Follow-Back (TLFB).

Because daily puff counts are non-negative count data and showed substantial overdispersion and a heavy right tail, we modeled them using a Bayesian Poisson-lognormal mixed-effects model. Unlike a standard Poisson model, which assumes that the conditional mean and variance of the outcome are equal, the Poisson-lognormal model allows the underlying daily puff rate to vary across observations through an additional normally distributed effect on the log-rate scale. This observation-level random effect accommodates excess day-to-day variability and extreme puff counts without requiring transformation of the observed count outcome.

Specifically, daily puff count for participant i on day t was modeled as:

$$Puff_{it} \sim Poisson(\lambda_{it})$$

with the expected puff rate defined as:

$$log(\lambda_{it}) = \beta_0 + \beta_{PE}\, PEcrave_{z,i} + \beta_{age}\, Age_{c,i} + \beta_{BMI}\, BMI_{c,i} + \beta_{sex}\, Sex_i + \beta_{day}\, DaySurvey_{c,it} + u_i + \varepsilon_{it}$$

Here, $u_i$ represents a participant-specific random intercept, assumed to follow a normal distribution with standard deviation $\sigma_{PID}$. This term accounts for stable between-participant differences in overall vaping frequency. The observation-level term $\varepsilon_{it}$ also follows a normal distribution, with standard deviation $\sigma_{obs}$, and captures residual day-to-day heterogeneity in puff rate beyond that expected under a standard Poisson distribution. Integrating over this normally distributed log-rate variation produces the Poisson-lognormal distribution.

Age and BMI were centered at their respective sample means, and sex was coded with female as the reference category. The day_survey variable was derived from Day_num modulo 7 and centered across observations to account for systematic variation in vaping across days of the weekly cycle. Analyses were restricted to observations with non-missing puff count, pattern expression, age, BMI, and sex, resulting in 58 participants and 1,407 person-days.

Weakly informative priors were assigned to the model parameters: Normal(0, 3) for the intercept, Normal(0, 2.5) for fixed-effect slopes, HalfNormal(1.5) for the participant-level random-intercept standard deviation, and HalfNormal(1.0) for the observation-level standard deviation. Posterior estimation was performed using the No-U-Turn Sampler (NUTS) with four chains, 1,000 tuning iterations and 1,000 posterior sampling iterations per chain, and a target acceptance rate of 0.95.

The association between vaping pattern expression and daily puff count was summarized using the posterior mean and standard deviation of the regression coefficient, the incidence rate ratio ($IRR = exp[\beta_{PE}]$) and its 95% highest-density interval (HDI), and the posterior probability that the association was positive, $P(\beta_{PE} > 0)$.

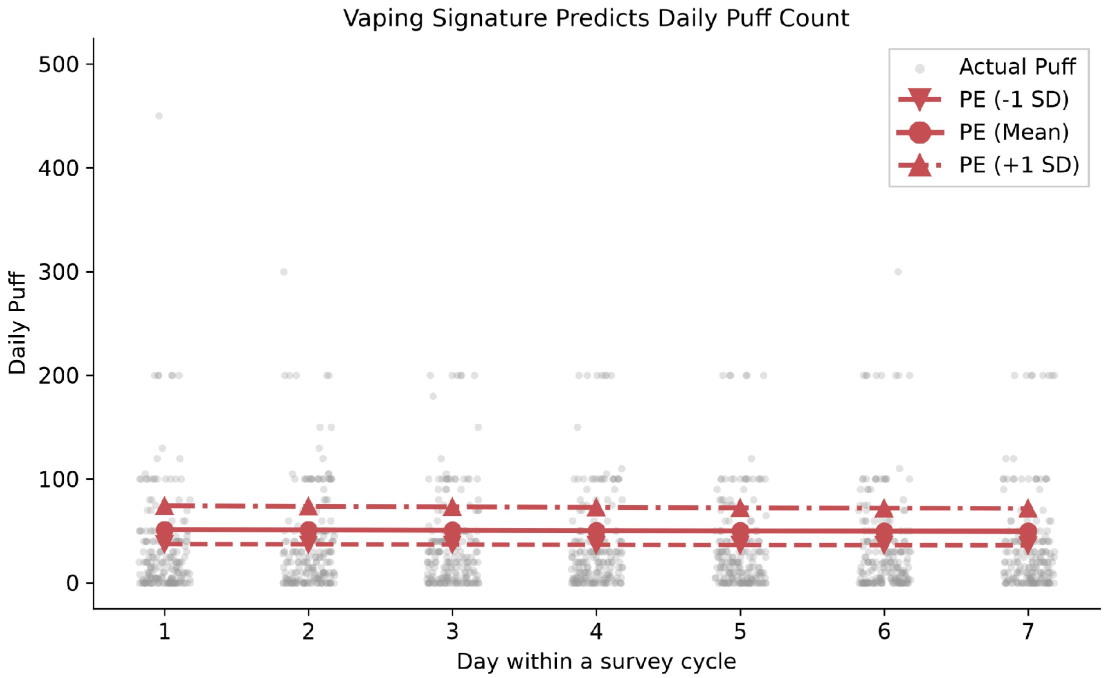


**Figure S1. Model-predicted versus observed daily e-cigarette puff counts, full range.** Same model-predicted daily puff counts by vaping-signature pattern expression (−1 SD, mean, +1 SD) and day within the weekly TLFB survey cycle as Figure 5B, shown here with the full range of observed daily puff counts (0–500) rather than the 0–100 cutoff used in the main text for visualization.

#### External dataset information

To standardize nicotine exposure and minimize severe withdrawal effects, participants were instructed to smoke immediately before arriving for the fMRI scan. During scanning, participants completed a cue-reactivity task consisting of two imaging runs, each lasting 430 s. Across the two runs, participants viewed four smoking-image blocks, four neutral-image blocks, four rest blocks, and craving-rating blocks. Smoking and neutral blocks each lasted 40 s and included eight images presented sequentially for 5 s each. Rest blocks consisted of a fixation cross presented for 20 s, and craving-rating blocks were presented for 10 s. Block order was counterbalanced across runs. Smoking blocks included images of cigarettes and people smoking, whereas neutral blocks included images of everyday objects, such as pencils and staplers, and people engaged in everyday activities, such as using a telephone. Participants were instructed to attend to the photographs presented on the screen. For fMRI data acquisition and preprocessing details, see Owens et al. (2018).

#### Low-level visual feature analysis

To assess whether low-level image properties differed across cue categories or covaried with normative craving, we quantified the low-level visual features of the 108 cue images (36 vaping, 36 smoking, and 36 food) presented in the cue-reactivity task. For each image, we computed 17 features spanning six domains: luminance (mean CIELAB lightness L* and its skewness), contrast (RMS contrast, i.e., the standard deviation of L* divided by 100, and a robust Michelson contrast based on the 1st and 99th percentiles of L*), color (mean CIELAB a* and b*, mean HSV saturation, and the colorfulness index of (Hasler & Süsstrunk, 2003)), edges (Sobel edge density and mean gradient magnitude), texture (grayscale Shannon entropy and gray-level co-occurrence matrix [GLCM] contrast, homogeneity, energy, and correlation, computed with 32 gray levels at a one-pixel distance and averaged over four orientations), and spatial frequency (the slope of the radially averaged amplitude spectrum of the Hann-windowed image in log–log coordinates and the proportion of power above half the Nyquist frequency). Edge, texture, and spatial-frequency features were computed on grayscale images resized to 256 × 256 pixels. All features were z-scored across images.

Redundant features were then removed with a two-step backward-elimination procedure applied to the z-scored features pooled across all 108 images. In Step 1 (pairwise redundancy), we computed the matrix of absolute Pearson correlations among the remaining features, identified the pair with the largest $|r|$, and, if it exceeded .85, dropped the member of that pair with the larger mean $|r|$ with all other remaining features (i.e., the more globally redundant feature). The correlation matrix was then recomputed and the step repeated until no pair exceeded $|r| = .85$. This step removed, in order, mean gradient magnitude ($|r| = .99$ with edge density), GLCM energy (.99 with entropy), GLCM homogeneity (.95 with edge density), edge density (.93 with GLCM contrast), high-frequency power (.90 with GLCM correlation), luminance skewness (.89 with mean luminance), and GLCM contrast (.86 with GLCM correlation). In Step 2 (multicollinearity), because pairwise correlations cannot detect a feature that is approximately a linear combination of several others, we computed for each remaining feature $j$ its variance inflation factor, $\text{VIF}_j = 1/(1 - R^2_j)$, where $R^2_j$ is the coefficient of determination from an ordinary least-squares regression (with intercept) of feature $j$ on all other remaining features. The feature with the largest VIF was dropped, the VIFs were recomputed on the reduced set, and the step was repeated until all VIFs were $\leq 5$. This step removed mean HSV saturation (VIF = 7.19) and then Michelson contrast (VIF = 5.10). The VIFs reported for the retained features are those of the final iteration, computed among the eight retained features only (range 1.51–3.28).

Each retained feature was then analyzed with a two-way ANOVA (category × craving level; Type III sums of squares with sum-to-zero contrasts), with partial eta-squared ($\eta_p^2$) as the effect size. To account for the dependence among images, category differences were followed up with paired $t$-tests across the 36 stimulus numbers (Holm-corrected within feature; effect size Cohen's $d_z$), and craving-level differences with Tukey HSD tests. We additionally tested linear trends across craving levels (coded 1–5), both averaged across categories and within each category, as well as the heterogeneity of trends across categories (trend × category). For each effect, $p$ values were corrected across features using the Benjamini–Hochberg false discovery rate (FDR; $q < .05$). Analyses were conducted in Python using statsmodels and SciPy.
